# Thermal acclimation in a non-migratory songbird occurs via changes to thermogenic capacity, but not conductance

**DOI:** 10.1101/2022.11.08.515677

**Authors:** Rena M. Schweizer, Abimael Romero, Bret W. Tobalske, Georgy Semenov, Matt D. Carling, Amber M. Rice, Scott A. Taylor, Zachary A. Cheviron

## Abstract

Thermoregulatory performance can be modified through changes in various subordinate traits, including thermal conductance, basal and summit metabolic rates, and body composition. We investigated physiological differences between black-capped chickadees (*Poecile atricapillus*) acclimated for six weeks to cold (−5°C) or control (25°C) environments (n = 7 per treatment) by measuring traits that affect thermal balance. We made repeated measurements of basal and summit metabolic rates via flow-through respirometry and body composition using quantitative magnetic resonance of live birds. At the end of the acclimation, we measured thermal conductance of the combined feathers and skins. Cold-acclimated birds had a higher summit metabolic rate, reflecting a greater capacity for endogenous heat generation, and an increased lean mass. However, birds did not alter their thermal conductance. These results suggest that, while birds can use multiple mechanisms to acclimate to their thermal environment, they use a subset of potential changes over varying timescales.

## Introduction

During winter, reductions in ambient temperature may present severe challenges to many animals. Animals exposed to the stress of seasonal cold temperatures may migrate elsewhere or stay and acclimate. For the latter, animals must endure decreased temperatures that often coincide with less daylight for foraging. For small non-hibernating species in particular, thermoregulatory performance can be a primary predictor of survival in cold environments (Fontanillas et al., 2005; Hayes and O’Connor, 1999; Lustick and Adams, 1977; Petit et al., 2017). For example, elevated thermogenic capacities are associated with enhanced survival probabilities in small endotherms that are native to cold climates (Fontanillas et al., 2005; Hayes and O’Connor, 1999; Petit et al., 2017).

Seasonal acclimatization allows endotherms to adjust to changes in temperature through physiological modifications that alter heat generation and/or retention, therefore enabling them to stay active during the winter. One strategy that birds can use to acclimate to winter temperatures is via increasing thermogenic capacity of the pectoralis and other muscles. Mechanisms for increasing thermogenesis could include increased muscle size for shivering, greater density of mitochondria, and/or changes to efficiency of mitochondrial activity (reviewed in Swanson, 2010). Additionally, birds can change conductive properties of the feathers and skin to diminish heat loss (Wolf and Walsberg, 2000). The relative importance of these two thermoregulatory strategies is not well understood, but is key to understanding how non-migratory birds survive harsh winters.

While a large body of work has advanced our understanding of how birds acclimatize in their summit metabolic rate (M_sum_, which represents total thermogenic capacity and may be measured as a cold-induced VO_2max_), and the subordinate traits that support it (e.g., (Barceló et al., 2017; Dubois et al., 2016; Petit et al., 2014; Swanson, 1991; Swanson, 1993; Swanson and Liknes, 2006), fewer studies have focused on acclimatization of body temperature regulation and the relative contributions of heat generation versus heat conservation mechanisms in that process. Recent research has demonstrated that dark-eyed juncos (*Junco hyemalis*) adjust their ability to thermoregulate over the course of a 9-week cold exposure via both increased metabolic heat production and decreased thermal conductance (Stager et al., 2020). Specifically, during a cold acclimation, juncos demonstrate increased heat generation (i.e., measured as M_sum_) after one week, yet show significantly increased heat retention (i.e., changes to conductance) only after nine weeks (Stager et al., 2020). This result suggests that juncos may meet specific thermal stresses of their environment by combining multiple responses that each respond to environmental variation over different timescales. However, the generality of this finding to other species of birds is unknown.

We investigated the patterns of seasonal acclimation in the black-capped chickadee (*Poecile atricapillus*), a small (9-14 g) passerine that is a year-round resident in temperate North America. Species within *Poecile* occupy one of the broadest altitudinal and latitudinal winter ranges of any passerine genus, and this distribution is shaped in part by thermoregulatory performance(Canterbury, 2002; Grossman and West, 1977; McQuillan and Rice, 2015; Olson et al., 2010; Swanson and Garland, 2009). High M_sum_ in birds is highly correlated with cold tolerance (Swanson, 2001; Swanson and Liknes, 2006). In free-living black-capped chickadees, elevated thermogenic capacities are associated with enhanced survival probabilities (Petit et al., 2017), and maximum cold-induced metabolic rate is correlated with thermogenic endurance (Swanson, 2001). A previous acclimation experiment of black-capped chickadees from Quebec, Canada, demonstrated that cold acclimation increased BMR (basal metabolic rate), M_sum_, and aerobic capacity of pectoralis muscle and liver tissues over a period of 28 days (4 weeks) (Milbergue et al., 2022). It remains unknown whether black-capped chickadees from other populations demonstrate similar increases in BMR and M_sum_, whether other traits that can influence thermoregulatory ability are affected, and what the time series is of acclimation in those and metabolic traits.

## Materials and Methods

### Collection and care of chickadees

We live-trapped 16 individuals from Boulder, CO, USA (40.0150° N, 105.2705° W; Colorado Parks and Wildlife Permit 1709964528, USFWS Permit MB06336A-1), using basket traps at feeders, on August 16, 2020, then drove them to Missoula, MT, USA, over 13 hours. All individuals were kept individually housed for both transport and the duration of the experiment. Upon initial arrival to Missoula, we inspected each bird for disease, assigned a sample identifier, and transferred it to a cage (76 cm x 45 cm x 45 cm). Birds were housed individually, and were given Factor 5 (Medpet, Benrose, South Africa) for five days for treatment against Mycoplasmosis, Canker (Trichomonas), Coccidiosis, Roundworm (Ascarids) and Hairworm (Capillaria). Birds had *ad libitum* access to food and water. Every day, individuals received a mix of sunflower seeds, ∼5-10 live mealworms, and ∼5-10 pine nuts, and every other day, ∼1 tsp of insect pattee (Orlux, Belgium) and ∼1 tsp of ground high-protein dog food. We supplemented water with vitamin drops (Wild Harvest, Blacksburg, VA, USA). For three weeks after arrival (August 17 to September 9, 2020), birds adjusted to captive conditions (23° C) and we aligned birds to a 12H:12H day:night schedule incrementally over that period. Birds were not observed to have undergone a molt while in captivity. All birds were maintained at this 12H:12H light:dark schedule for the remainder of the experiment. Two birds died during the first week of captivity. Sex was determined by identification of gonads after euthanasia (8 females, 6 males). All animal care was approved by the University of Montana IACUC (AUP 043-19).

### Acclimation trials and metabolic assays

After the initial period of adjustment to captive conditions, we randomly assigned individuals to one of two 6-week acclimation treatments of either control (23°C; n=7 birds) or cold (−5°C; n=7 birds) temperatures. We measured birds at week 0 (prior to placement in experimental treatment), week 3, and week 6 of the acclimation (3, 6, and 9 weeks after capture, respectively), as follows. To measure basal metabolic rate (BMR), we removed food from cages at least 2 hours prior to ensure birds were postabsorptive prior to measurement (as in (Olson et al., 2010), and used a Sable Systems flow-through respirometry system. One to two birds from each treatment were measured each day, with trials starting at approximately 17:00 h. Prior to the BMR measurement, we weighed each bird using a portable balance (Fisher Scientific). Each bird was placed in a 1L volume Nalgene bottle chamber outfitted with input and output connection for air flow into and out of the chamber, and the bird was placed inside a dark temperature-controlled chamber (Sable Systems Pelt Cabinet with Pelt-5 Temperature Controller, North Las Vegas, NV, USA) held at ∼32°C. This temperature is within the thermoneutral zone of chickadees (Olson et al., 2010; Rising and Hudson, 1974). We dried ambient air using Drierite, then used an FB8 mass flow meter (Sable Systems), to control air flow into each chamber at ∼500 mL/min. For each sample or baseline chamber, we manually subsampled output airflow at 100 - 150 ml/min using barrel syringes, dried air with Drierite, scrubbed CO_2_ using Ascarite, then re-dried air with Drierite prior to measuring O_2_ using a FoxBox (Sable Systems). We let the birds adjust and acclimate to the respirometry chambers for ∼15 minutes, then we started measuring with an empty baseline chamber of 15 min, followed by measuring the bird chamber for 15 min. Bird and baseline chambers were all 1L Nalgene containers. Three individually-housed birds were measured concurrently by cycling through each bird three times, with a 15 min baseline between each measurement, for a total measurement period of 45 min per bird over 3 hours. Each day, before the respirometry experiments, we spanned the FoxBox using ambient air (20.95% O_2_). At the end of the BMR trials, birds were returned to their acclimation treatment and were provided food.

The morning after each BMR trial, we measured M_sum_, or the cold-induced maximum metabolic rate (VO_2_max), for each bird as follows. Starting at approximately 9:00 h, we weighed each bird and placed it in a chamber within a temperature cabinet kept at ∼-5°C. Incurrent air to the chamber was a mix of 21% oxygen and 79% helium, with flow rates and mixing regulated using a mass flow controller (Alicat Scientific, Tucson, AZ, USA) with a flow rate of ∼750 ml/min. Given the higher conductance of heliox over that of ambient air, the use of heliox during measurement of M_sum_ enables eliciting a response to cold stress at higher ambient temperatures than without heliox, resulting in less potential for damage to the organism (Rosenmann and Morrison, 1974; Swanson, 2010). After a 5 minute baseline measurement, we measured each bird’s oxygen consumption for a maximum 30 minute period or until they became hypothermic (visible by a peak of oxygen consumption followed by a sharp decline). Before and after each trial, we measured the bird’s body temperature using a probe inserted into the cloaca; birds with a body temperature at or below 37°C were considered hypothermic. At week 3 and week 6, control birds had their M_sum_ measured in a temperature cabinet at -5°C, as at week 0, while birds that were in the cold treatment had their M_sum_ measured in a temperature cabinet (Accucold VLT65, Bronx, NY, USA) maintained at ∼-20°C. We reasoned that if birds in the control group are hypothermic at the end of the -5°C trial, they likely would have been as cold challenged as the cold acclimated birds at -20°C, given that hypothermia suggests the inability to regulate body temperature. These conditions were similar to those implemented previously (Stager et al., 2020). At week 3, 12 of 14 birds had a body temperature ≤37°C (indicating hypothermia; (Stager et al., 2020; Swanson et al., 2014). Two birds did not have reliable body temperature estimates, but their oxygen traces indicated M_sum_ was reached so they were included in analysis. At week 6, all birds had a body temperature ≤37°C after the M_sum_ trial. A single researcher (R.M.S.) performed all metabolic assays.

Following Stager et al. (2020) and the custom R scripts therein (https://github.com/Mstager/batch_processing_Expedata_files), we corrected for fluctuations in baseline measurements using a linear correction, then, following (Lighton, 2008), calculated BMR as the lowest oxygen consumption rate (ml O_2_/min) averaged over a 10-min period. After baseline corrections, we calculated M_sum_ as the highest oxygen consumption (ml O_2_/min) over a 5-min period.

### Physiological phenotyping

At week 0, week 3, and week 6 of the acclimation experiments, we measured body fat and lean mass of each bird using an EchoMRI quantitative magnetic resonance (QMR) system (E26-281-BHlab, Houston, TX, USA) after M_sum_ trials. We measured three standards at the beginning and end of each day’s QMR measurements, then calculated a standard curve and adjusted the sample values accordingly using linear regression in R. After week 6 of the acclimation experiment, we euthanized birds via thoracic compression and dissected out organs, determining sex via identification of the gonads. We measured hemoglobin concentration using a HemoCue Hb 201 (Brea, CA, USA) and, after spinning down blood-filled capillary tubes for 5 min in a centrifuge, measured hematocrit as the relative proportion of packed red blood cells using digital calipers. We filled each body cavity with a damp paper towel, then froze the skins at -20°C until the conductance assays were performed.

### Conductance assays

To measure skin and plumage thermal conductance, we quantified the power input required to maintain a constant internal temperature of 39 °C as a relative measure of heat loss to ambient temperature. Before conducting trials, each skin was thawed and any remaining adipose or muscle tissue was removed from the body cavity. Using corn meal we brushed feathers to remove any excess oils. We then created an epoxy mold (fit to the internal cavity of a ∼9 g chickadee) composed of a thermocouple and nichrome wire contained in PC-Marine Epoxy Putty (Allentown, PA, USA). Using needle and thread, we stitched the skins around the mold within the thoracic and abdominal cavity. Skins were then placed in an upright position hanging by threads attached to external nares with wings tucked to sides. We monitored temperature of the mold via a thermocouple, and measured the power input (milliwatts, mW) necessary to maintain the epoxy mold at a constant temperature using a voltage logger (Omega OM-CP-Quadvolt, Norwalk, CT, USA), an amperage logger (Omega OM-CP-Process 101A-3A) and a temperature controller (Omega CNI1622-C24-DC). A 12 V DC battery was used to power the board as well as to heat the epoxy mold.

We performed conductance trials in a small room with limited airflow and a relatively constant ambient room temperature (mean±s.d. = 23.7 ±0.7°C). Epoxy molds were heated to 39°C and power was required to increase temperature of the thermocouple in the mold when it fell below 37°C. Throughout a 20 minute period both amperage and Voltage were recorded at every second to measure the power input required to maintain an internal temperature of 39°C. We calculated power input by taking the mean of Volts*amps over the 20 minute interval for a sample size of n=14. Trials were conducted by a single person (A.R.) blind to each bird’s treatment. We did not find a significant effect of variation in room temperature on average power (p=0.872), and only a marginally significant effect of sampling date (0.08545) using linear regression models. When modeling the effect of treatment on average power, we compared a model with sampling date to a model without, and the model without sampling date had a lower AICc (Avg. Power ∼ Treatment + Date: AICc=193.3; Avg. Power ∼ Treatment: AICc=162.3).

### Statistical analyses

We performed all statistical analyses in R, and the code and R notebook are available on Github (https://github.com/renaschweizer/chickadee_thermal_acclimation_paper). Similar analyses for all phenotypic traits were performed as follows. First, we quantified the effects of acclimation on phenotypic traits with linear mixed-effects models using the Lme4 package in R (Bates et al., 2015). For BMR and M_sum_ we tested for an effect of a treatment*timepoint interaction, including body mass (M_b_) as a covariate and bird ID as a random effect (e.g., phenotype ∼ mass + treatment*timepoint + (1| bird ID)). For M_b_, we included the treatment*timepoint interaction with bird ID as a random effect. We used ANOVA tests to assess significance, and compared models using the AICc implementation in MuMIn (Barton, 2010). To quantify the effects of treatment acclimation on conductance, we used a standard linear regression model with random effects removed (Lm function in R). We used post-hoc tests to find pairwise differences of treatment*timepoint on phenotypes with significant treatment or timepoint effects using the emmeans package in R and a Benjamini-Hochberg correction for multiple testing. We also tested treatment and timepoint effects on hemoglobin, percent red blood cells (hematocrit), and heart mass. Finally, we tested for significant pairwise associations between phenotypes using Pearson correlation (cor.test in base R).

## Results and Discussion

### Cold acclimation increases thermogenic capacity

After adjusting to captivity, but prior to experimental acclimation, chickadees did not differ by treatment in their BMR (p=0.6704) nor their M_sum_ (p=0.8303) values (Table S1; Table S2), demonstrating that individuals did not differ in this trait prior to temperature acclimation. Birds gained body mass over the experimental acclimation (Figure S1), with timepoint and a treatment*timepoint interaction having significant effects (Table 1). BMR and M_sum_ were highly and significantly correlated (r^2^: 0.325; p-value: 0.036), but were not correlated once mass-corrected (r^2^: 0.099; p-value: 0.534).

**Table 1.**
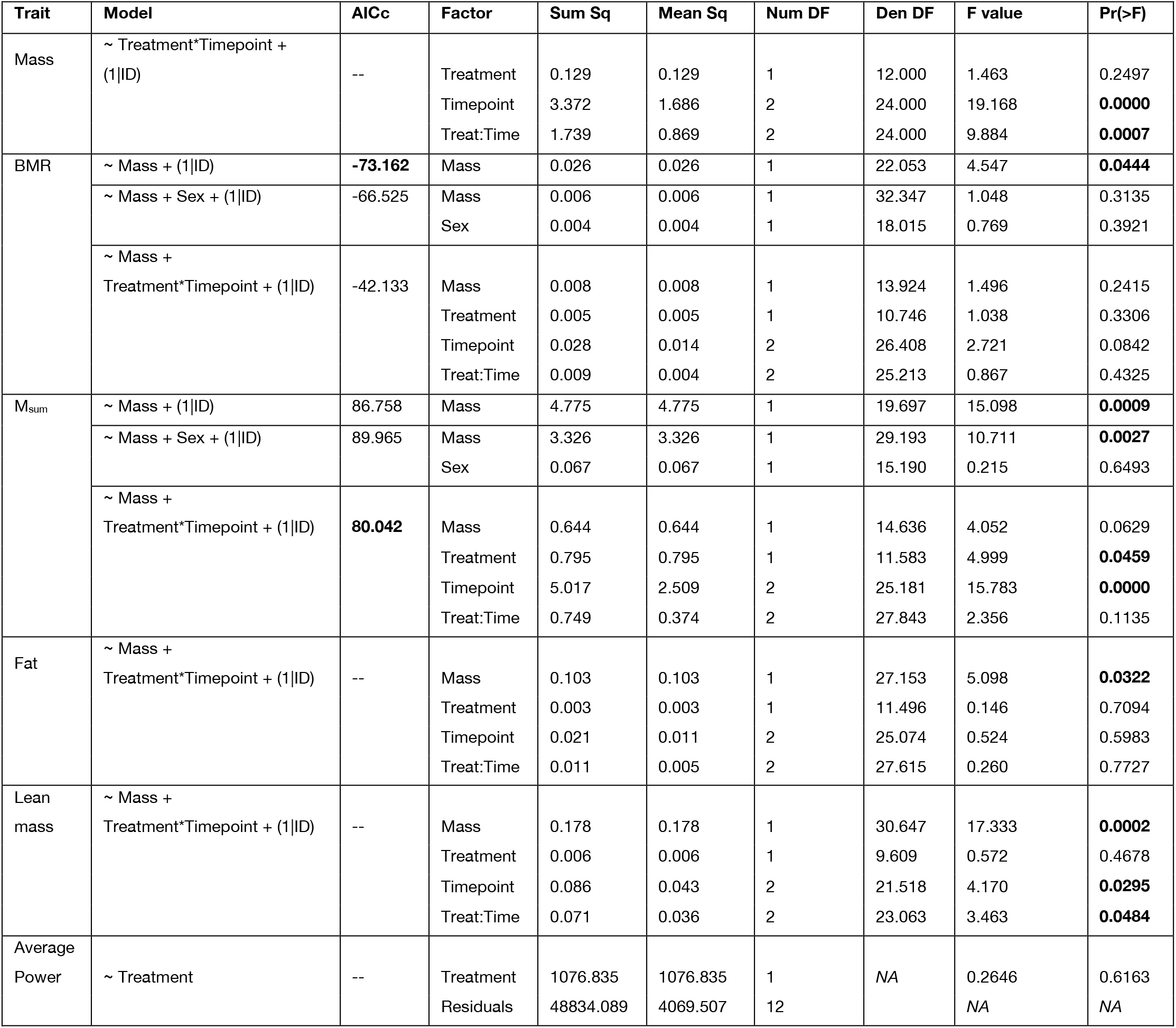
Results of linear models testing the effects of mass, treatment, timepoint, and the treatment x timepoint interaction (Treat:Time) on various physiological measurements. For BMR and M_sum_, multiple models were tested and the corresponding AICc values are reported. The table reports results of reduced linear mixed effect models. NumDF and DenDF are the numerator and denominator degrees of freedom, respectively. P-values ≤0.05 are bolded.

| Trait | Model | AICc | Factor | Sum Sq | Mean Sq | Num DF | Den DF | F value | Pr(>F) |
| --- | --- | --- | --- | --- | --- | --- | --- | --- | --- |
| Mass | ~ Treatment*Timepoint + (1\|ID) | -- | Treatment | 0.129 | 0.129 | 1 | 12.000 | 1.463 | 0.2497 |
|  |  |  | Timepoint | 3.372 | 1.686 | 2 | 24.000 | 19.168 | **0.0000** |
|  |  |  | Treat:Time | 1.739 | 0.869 | 2 | 24.000 | 9.884 | **0.0007** |
| BMR | ~ Mass + (1\|ID) | **-73.162** | Mass | 0.026 | 0.026 | 1 | 22.053 | 4.547 | **0.0444** |
|  | ~ Mass + Sex + (1\|ID) | -66.525 | Mass | 0.006 | 0.006 | 1 | 32.347 | 1.048 | 0.3135 |
|  |  |  | Sex | 0.004 | 0.004 | 1 | 18.015 | 0.769 | 0.3921 |
|  | ~ Mass + Treatment*Timepoint + (1\|ID) | -42.133 | Mass | 0.008 | 0.008 | 1 | 13.924 | 1.496 | 0.2415 |
|  |  |  | Treatment | 0.005 | 0.005 | 1 | 10.746 | 1.038 | 0.3306 |
|  |  |  | Timepoint | 0.028 | 0.014 | 2 | 26.408 | 2.721 | 0.0842 |
|  |  |  | Treat:Time | 0.009 | 0.004 | 2 | 25.213 | 0.867 | 0.4325 |
| $M_{\text{sum}}$ | ~ Mass + (1\|ID) | 86.758 | Mass | 4.775 | 4.775 | 1 | 19.697 | 15.098 | **0.0009** |
|  | ~ Mass + Sex + (1\|ID) | 89.965 | Mass | 3.326 | 3.326 | 1 | 29.193 | 10.711 | **0.0027** |
|  |  |  | Sex | 0.067 | 0.067 | 1 | 15.190 | 0.215 | 0.6493 |
|  | ~ Mass + Treatment*Timepoint + (1\|ID) | **80.042** | Mass | 0.644 | 0.644 | 1 | 14.636 | 4.052 | 0.0629 |
|  |  |  | Treatment | 0.795 | 0.795 | 1 | 11.583 | 4.999 | **0.0459** |
|  |  |  | Timepoint | 5.017 | 2.509 | 2 | 25.181 | 15.783 | **0.0000** |
|  |  |  | Treat:Time | 0.749 | 0.374 | 2 | 27.843 | 2.356 | 0.1135 |
| Fat | ~ Mass + Treatment*Timepoint + (1\|ID) | -- | Mass | 0.103 | 0.103 | 1 | 27.153 | 5.098 | **0.0322** |
|  |  |  | Treatment | 0.003 | 0.003 | 1 | 11.496 | 0.146 | 0.7094 |
|  |  |  | Timepoint | 0.021 | 0.011 | 2 | 25.074 | 0.524 | 0.5983 |
|  |  |  | Treat:Time | 0.011 | 0.005 | 2 | 27.615 | 0.260 | 0.7727 |
| Lean mass | ~ Mass + Treatment*Timepoint + (1\|ID) | -- | Mass | 0.178 | 0.178 | 1 | 30.647 | 17.333 | **0.0002** |
|  |  |  | Treatment | 0.006 | 0.006 | 1 | 9.609 | 0.572 | 0.4678 |
|  |  |  | Timepoint | 0.086 | 0.043 | 2 | 21.518 | 4.170 | **0.0295** |
|  |  |  | Treat:Time | 0.071 | 0.036 | 2 | 23.063 | 3.463 | **0.0484** |
| Average Power | ~ Treatment | -- | Treatment | 1076.835 | 1076.835 | 1 | NA | 0.2646 | 0.6163 |
|  |  |  | Residuals | 48834.089 | 4069.507 | 12 | NA | NA | NA |

BMR and M_sum_ were both significantly affected by M_b_ but not by sex (Table 1). BMR was significantly affected by and best explained by M_b_ (p-value: 0.044; BMR ∼ M_b_ + (1|ID)); in a more complex model (BMR ∼ M_b_ + Treatment*Timepoint + (1|ID)), we found no significant effect of M_b_, treatment, timepoint, or interaction on BMR (Figure 1A; Table 1; Figure S2). However, when we examined the week 6 BMR data separately, we did find a significant effect of treatment (p=0.048) and mass (p=0.028), with a higher BMR in cold birds (Table S4).

**Figure 1.**
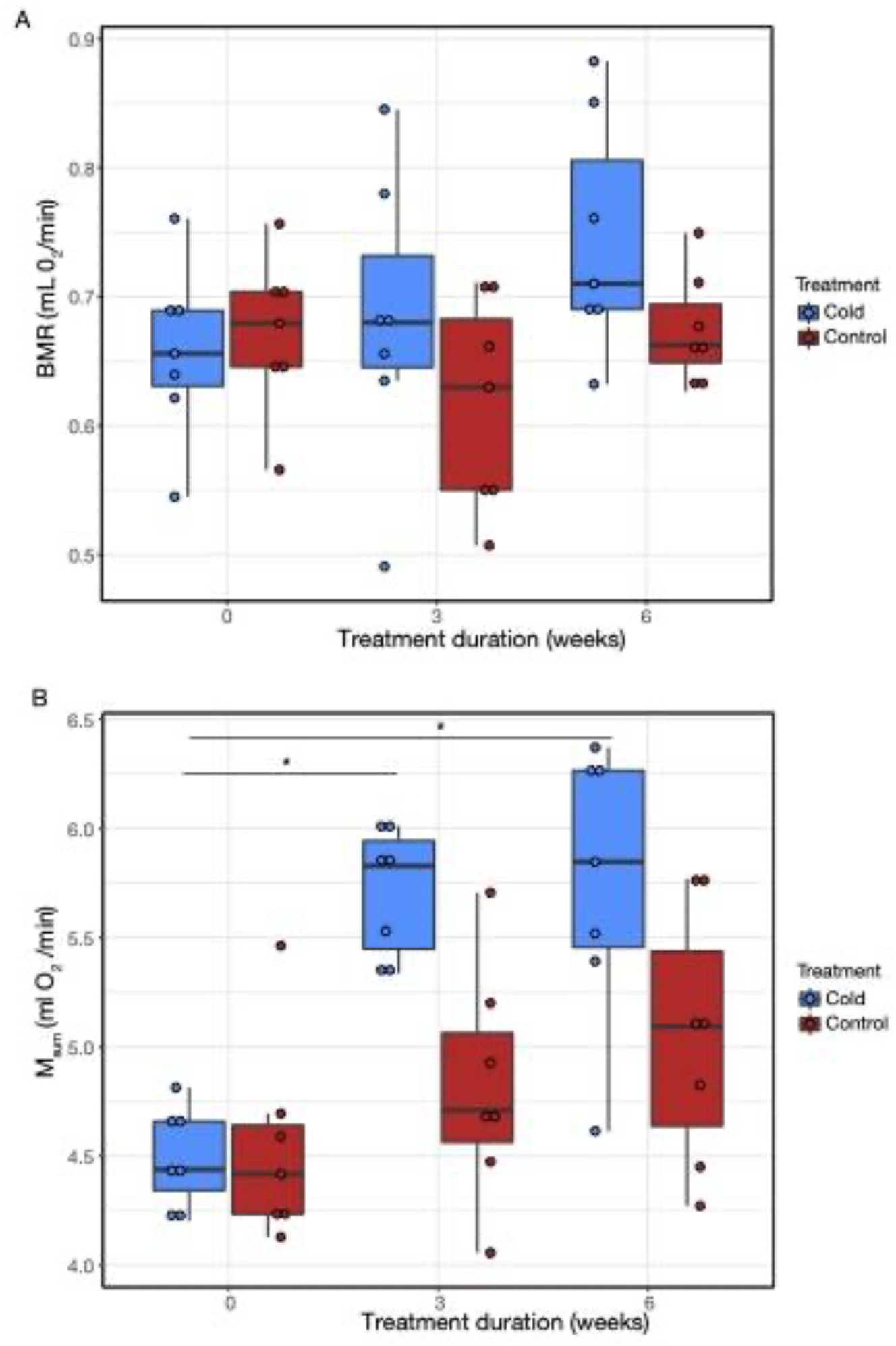
Repeated measures of BMR and M_sum_ for control (23°C, n=7) and cold (−5°C, n=7) experimental treatments. A) Over the 6 week acclimation, there was no significant effect of treatment or timepoint on BMR. B) Chickadees showed significant effects of treatment (p<0.05) and time point (p<0.01) on M_sum_, with significant changes in cold-acclimated M_sum_ from T_0wk_ to T_3wk_ (p=0.0010) and from T_0wk_ to T_6wk_ (p=0.0010), and in control-acclimated M_sum_ from T_0wk_ to T_6wk_ (p=0.0259).

We found significant effects on M_sum_ of treatment (p=0.046) and time point (p=3.607e-05) (Figure 1B), but no effect of M_b_ (0.062)(Table 1; Figure S3). The best fit model for M_sum_ included M_b_, treatment*timepoint, and bird ID, but not sex (Table 1). Cold treatment birds increased their M_sum_ from T_0wk_ to T_3wk_ (p=0.001) and from T_0wk_ to T_6wk_ (p=0.001), but not from T_3wk_ to T_6wk_ (p=0.986)(Table S3). Control birds also increased their M_sum_ from T_0wk_ to T_3wk_ (p=0.001) and from T_0wk_ to T_6wk_ (p=0.026, but not from T_3wk_ to T_6wk_ (p=0.275)(Table S3). At both T_3wk_ and T_6wk_, the mean M_sum_ was significantly different between treatment groups (p=0.003 and p=0.022, respectively), with cold-acclimated birds having an M_sum_ elevated by ∼14-18% (mean values for T_3wk_: M_sum, cold_=5.708, M_sum, control_=4.818; mean values for T_6wk_: M_sum, cold_=5.753, M_sum, control_=5.041). M_b_ had a significant effect on M_sum_ at T_6wk_ but not T_3wk_ (Table S4).

Over our acclimation experiment, black-capped chickadees increased M_sum_ in the cold, but BMR did not increase significantly (Figure 1), except when analyzed only with week 6 data (Table S4). BMR and M_sum_ are likely to be functionally linked for a few potential reasons. The most common explanation is that the pectoralis muscles tend to grow with cold exposure to support higher shivering capacity (see discussion on lean mass, below), and it follows that muscle hypertrophy may necessitate higher energetic costs that are reflected in BMR (Kersten and Piersma, 1987; McNab, 1994). In some contexts, cold-induced changes in BMR are associated with an increase in food intake and the resulting hypertrophy of digestive organs (Barceló et al., 2017), however, these effects are not universal (Petit et al., 2014; Vézina et al., 2017). Previous studies also suggest that in some birds, BMR is lower in winter compared to summer (Petit and Vézina, 2014; Saarela et al., 1995; Smit and McKechnie, 2010) pointing to a disconnect between BMR and thermoregulatory requirements.

Contrary to some previous findings (Dubois et al., 2016; Fristoe et al., 2015; Liknes et al., 2002; Milbergue et al., 2022; Nilsson and Nilsson, 2016) but see Smit et al. 2010), cold treatment was not a significant factor explaining BMR (Figure 1A). Cold-acclimated birds did demonstrate a higher mean BMR after 6 weeks than control birds (cold: 0.745 mL O_2_/min; control: 0.675 O_2_/min), even after accounting for mass differences between the two groups (cold: 0.059 mL O_2_/min/g; control: 0.058 O_2_/min/g). However, this difference was significant only when the data from week 6 was compared, and was not significant in the full model with treatment*timepoint (Table 1; Table S3). It is possible that the -5°C cold treatment was not harsh enough to cause a significant change in BMR. It is also possible that BMR and M_sum_ are not functionally linked in this population of black-capped chickadees. In black-capped chickadees from Quebec, Canada, Petit et al. (2014) found minimal, non-significant seasonal changes in BMR, but seasonal increases in M_sum_. Yet, other studies of black-capped chickadees have identified significant cold acclimatization (Cooper and Swanson, 1994; Swanson and Liknes, 2006) and acclimation effects (Milbergue et al., 2018; Milbergue et al., 2022).

We did not measure the masses of digestive organs or food intake so we are unable to assess whether cold-acclimation affected these traits. If changes in BMR are uncoupled from M_sum_, they may be tied to fitness in unmeasured ways, or linked to other behavioral (e.g., energy expenditure) or metabolic traits. For example, Milbergue et al. (2022) found that, in cold-acclimated black-capped chickadees, heightened BMR and M_sum_ were correlated with divergent patterns of mitochondrial metabolism in different tissues (i.e., with complex I activity in the liver and with mitochondrial proton leak in the muscle, respectively).

Our M_sum_ results are consistent with previous studies demonstrating an increased ability to tolerate cold in winter-acclimatized birds relative to those sampled in the summer (Cooper and Swanson, 1994) and in cold-acclimated birds relative to control (Milbergue et al. 2022). M_sum_ is a proxy for the maximum thermogenic capacity of an endotherm, and as a result, is positively correlated with cold tolerance (i.e., endurance under cold stress; Swanson 2001; Swanson et al., 2006; Swanson et al., 2009; Barceló et al. 2017). Increases in thermogenic capacity underscore the importance of internal heat generation as chickadees acclimate to colder temperatures.

### Cold acclimation increases lean mass, but not fat mass

We did not find an effect of treatment or timepoint on total fat mass over the acclimation period (Figure S4; Table 1). However, upon examination of lean mass, there was a significant effect of timepoint (p=0.030), and treatment*timepoint (p=0.048) (Figure 2A; Table1). As expected, body mass had significant effects on both total fat mass and lean mass (Table 1), and lean mass significantly increased over time with cold acclimation (Figure 2A). Within cold-acclimated birds, lean mass was significantly different from T_3wk_ to T_6wk_ (p=0.036), but there were no significant pairwise differences within the control birds. Throughout the acclimation period, overall mass increased (Figure S1). While chickadees did not alter fat mass over the experimental period, there was a non-significant trend towards increased total fat mass in cold-acclimated birds (Figure S4).

**Figure 2.**
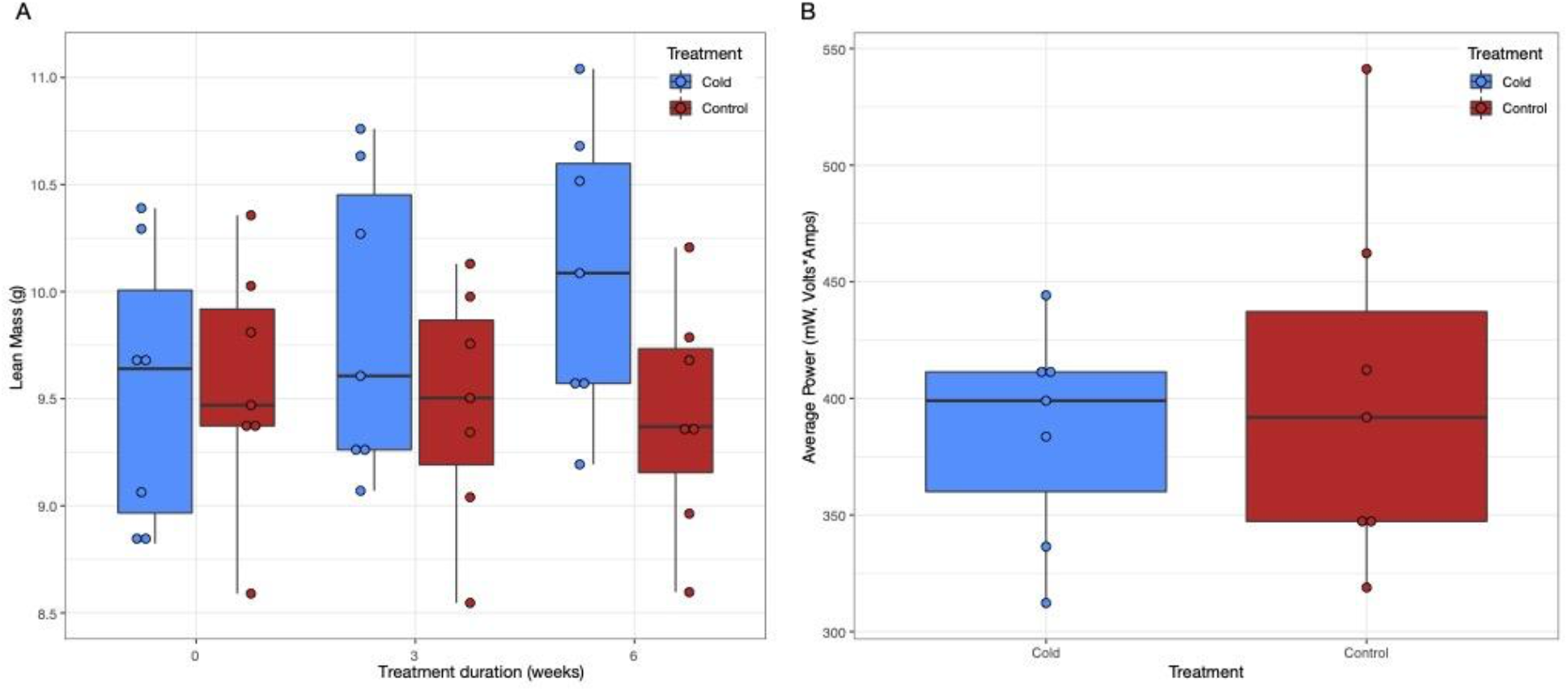
Lean mass and conductance over experimental acclimation. A) Lean mass was significantly affected by timepoint (p=0.0295) and timepoint*treatment (p=0.0484), and significantly increased in cold-acclimated birds from T_3wk_ to T_6wk_ (p=0.0359). B) Average power measured between birds from cold and control treatments. Power was not significantly affected by treatment (p>0.05).

The lack of significant increases in fat mass is in contrast to previous studies on free-living dark-eyed juncos and black-capped chickadees that found increases in winter fat stores (Chaplin, 1974; Swanson, 1991). While one of the functions of fat is to provide fuel stores for wintering birds, studies have demonstrated that fat mass may not directly influence thermogenic capacity (Marsh and Dawson 1989). Because M_sum_ in birds is largely the result of active shivering by skeletal muscles, the flight muscles (pectoralis) are considered to be the primary thermogenic organ (Marsh and Dawson 1989; Swanson et al., 2013). Seasonal flexibility of M_sum_ should thus be rooted in seasonal flexibility of the pectoralis. An important mechanism for increasing organismal thermogenic capacity in winter is indeed to increase the size of the pectoralis, and has been observed in a variety of bird species (Swanson and Vézina, 2015).

While we did find an increase in lean mass, the time course of lean mass increase was delayed compared to the increase in M_sum_, suggesting that improvements in M_sum_ cannot be solely attributable to pectoralis growth (Figure 2A). Stager et al. (2020) similarly found in dark-eyed juncos that thermogenic mechanisms change over different timescales, given that temperature regulation and resistance to hypothermia continued to improve in the absence of improvements in M_sum_. Although pectoralis muscle size is a prominent predictor of M_sum_ in birds (Swanson, 2010), other factors that influence aerobic capacity of muscle fibers could explain improvements in M_sum_ in the absence of changes to lean mass. For example, changes in mitochondrial densities, metabolic enzyme activities, or blood supply to respiring tissues have all been shown to influence metabolic capacity (Suarez, 1998). In cold-acclimated black-capped chickadees, specifically, elevated M_sum_ has been positively correlated with mitochondrial proton leak (Milbergue et al., 2022), which has been suggested as a heat-generating mechanism.

Changes in these factors could alter metabolic intensity without concurrent changes in muscle mass (Milbergue et al., 2018; Swanson, 2010). While we did measure hemoglobin concentration, hematocrit, and heart mass, none of these traits differed significantly between treatment conditions (Table S4). Other studies have also found increases in mass of body muscles (pectoralis, legs, supracoracoideus, skeletal) and cardiopulmonary organs as well as hematocrit levels during winter (Petit et al., 2014; Swanson, 1991; Swanson, 2010). The overall increase in lean mass (including the pectoralis muscle) are consistent contributors to the winter phenotype of small birds in cold climates (Liknes and Swanson, 2011; Petit et al., 2013; Vézina et al., 2011).

### Chickadees do not modify their thermal conductance in response to cold treatment

Despite the expectation that changes in heat retention could improve thermoregulation in the cold, we found that cold acclimation did not significantly affect thermal conductance (Figure 2B; Table 1). This suggests that black-capped chickadees do not simultaneously alter physiological processes that improve both capacity to generate (Figure 1) and retain (Figure 2B) heat, at least over the timescales studied here.

While our study shows a lack of significant difference in thermal conductance between birds acclimated to -5°C and birds acclimated to 23°C, it is important to note previous studies that have found increase in conductance for birds acclimated to summer temperatures (Swanson, 1991) as well as decreasing conductance for cold-acclimated birds (Stager et al., 2020). One possible explanation could be simply that our experimental trial lengths were not long enough to show changes. For example, Stager et al. (2020) found that cold-acclimated dark-eyed juncos showed reduced conductance only at 9 weeks (and not at weeks 1, 2, 3, or 6). However, Stager noted that the condition of birds at 9 weeks suggested overall poor health, perhaps as a result of captive conditions (pers. comm.), that likely increased stress in captivity (Dickens et al., 2009; Fischer and Romero, 2019). This led us to end the experiment at 6 weeks.

An additional important factor to consider is that conductance may respond to different seasonal cues. Specifically, large changes to conductance in birds may require a molt, which is often triggered by changes in photoperiod (reviewed in (Dawson et al., 2001)). In a study focusing on red knots, an arctic shorebird, seasonal changes in thermal conductance were likely due at least in part to changes in insulative feathers (Piersma et al., 1995). Additionally, Swanson (1991) measured the dry mass of contour plumage in dark-eyed juncos, where the contour plumage was an index of insulation, and found the mass increased by 37% in birds captured during the winter. While temperature and photoperiod covary as seasonal cues and the full seasonal phenotype may be the outcome of response to one or the other cue, in our experiment we only modified temperature and kept photoperiod constant. Black-capped chickadees typically molt once a year in the late summer, so our individual birds were collected after molting, leaving them with little ability to modify feathers during the experiment.

However, there are a variety of other factors that might influence thermal conductance in birds, such as total surface area, skin composition, and behavior (Wolf and Walsberg, 2000). Skin inherently plays a role in thermoregulation, and studies have pointed to variation in plumage densities and skin surface area being related to enhanced cold tolerance (Swanson, 1993). Studies have shown that plastic changes in lipid composition of the outer layer of the epidermis is important for avoiding water loss in desert birds (Muñoz-Garcia and Williams, 2008; Muñoz-Garcia et al., 2008). For wintering birds, similar plasticity in skin composition could affect heat retention for enduring cold temperatures. Additionally, birds may thermoregulate through behaviors such as clustering with conspecifics (McKechnie and Lovegrove, 2001) or posturing (Ferretti et al., 2019).

### Mechanisms of thermoregulation in black-capped chickadees

Our results suggest that, in response to cold, black-capped chickadees modify their thermoregulatory ability through changes in internal heat generation rather than changes in heat retention, with a few caveats. The 6-week acclimation period may not have been long enough to capture additional changes in thermal conductance, or the birds may have arrived in Montana after molting in the wild. Regardless, our results indicate that chickadees may be able to respond to short-term bouts of thermal stress by increasing their M_sum_ rather than changing their conductance. Cold snaps might therefore require a greater energetic investment to offset if they are solely dealt with via changes to metabolism. To better understand the underlying mechanisms driving thermoregulation in birds, it would be beneficial to study both the behavioral and physiological aspects of the avian response to cold throughout the acclimation period. Further studies could also shed light on specific physiological changes that contribute to improvements in whole-organism performance.

## Acknowledgements

We thank K. Wilsterman for advice on data analysis and M. Stager for help with input on experimental design and data processing. We are grateful to members of the Cheviron lab for feedback and for help with bird care.

## Competing interests

The authors declare that no competing interests exist.

Rena M. Schweizer^1,5^, Abimael Romero^1^, Bret W. Tobalske^1^, Georgy Semenov^2^, Matt Carling^3^, Amber M. Rice^4^, Scott A. Taylor^2^, Zachary A. Cheviron^1^

## Author contributions

Conceptualization: RMS, SAT, ZAC, MDC

Methodology: RMS, BWT, GS, SAT, ZAC

Formal analysis: RMS, AR

Resources: BWT, SAT, ZAC

Data curation: RMS, AR

Writing - original draft: RMS, AR

Writing - review & editing: BWT, SAT, AMR, GS, ZAC, MDC

Funding acquisition: BWT, SAT, ZAC, MDC

## Data Availability

Metadata compiled for each individual is available in Table S1 and provided as individual files with a R data analysis script on R.M.S.’s Github page (https://github.com/renaschweizer/chickadee_thermal_acclimation_paper). Raw respirometry, QMR, and conductance data are available upon request. Scripts for processing raw respirometry data are similar to those available on Github (https://github.com/Mstager/batch_processing_Expedata_files).

## Supplemental Tables

**Table S1**. Meta data for repeated experimental measures of mass, BMR, SMR, and associated phenotypes for control (23°C, n=7) and cold (−5°C, n=7) experimental treatments.

**Table S2**. Results of linear models testing the pre-experimental treatment effects of mass and treatment on BMR and M_sum_.

**Table S3**. Results of linear models testing the effects of mass, treatment, timepoint, and the treatment x timepoint interaction (Treat:Time) on various physiological measurements. For BMR and M_sum_, multiple models were tested and the corresponding AICc values are reported. Extended version of Table 1 showing contrasts.

**Table S4**. Results of linear models testing the end-of-experiment treatment effects of mass and treatment on BMR, M_sum_, hemoglobin, percent RBC, and heart mass.

## Supplemental Figures

**Figure S1**. Repeated measures of body mass for control (23°C, n=7) and cold (−5°C, n=7) experimental treatments. Body mass was measured in grams prior to BMR measurement.

**Figure S2**. Repeated measures of BMR (mass-corrected) for control (23°C, n=7) and cold (−5°C, n=7) experimental treatments.

**Figure S3**. Repeated measures of M_sum_ (mass-corrected) for control (23°C, n=7) and cold (−5°C, n=7) experimental treatments.

**Figure S4**. Repeated measures of total fat mass (g) for control (23°C, n=7) and cold (−5°C, n=7) experimental treatments.

